# Analyzing functional heterogeneity of effector cells for enhanced adoptive cell therapy applications

**DOI:** 10.1101/2024.05.27.595942

**Authors:** Anne-Christine Kiel Rasmussen, Thomas Morgan Hulen, David Leander Petersen, Mette Juul Jacobsen, Marie Just Mikkelsen, Özcan Met, Marco Donia, Christopher Aled Chamberlain, Peter Mouritzen

## Abstract

Cellular effector function assays traditionally rely on bulk cell populations that mask complex heterogeneity and rare subpopulations. The Xdrop® droplet technology facilitates high-throughput compartmentalization of viable single cells or single-cell pairs in double-emulsion droplets, enabling the study of single cells or cell-cell interactions at an individual level. Effector cell molecule secretion and target cell killing can be evaluated independently or in combination. Compatibility with a wide range of commercial assay reagents allows for single-cell level readouts using common laboratory techniques such as flow cytometry or microscopy. Moreover, individual cells of interest can be viably isolated for further investigation or expansion. Here we demonstrate the application of the double-emulsion droplet technology with a range of cell types commonly utilized for adoptive cell therapy of cancer: peripheral blood mononuclear cells, natural killer cells, tumor-infiltrating lymphocytes, and chimeric antigen receptor T cells. Single-cell compartmentalization offers unparalleled resolution, serving as a valuable tool for advancing the development and understanding of cellular therapy products.

## Background

Adoptive cell therapies (ACT) have demonstrated impressive curative effects against specific cancer types, as illustrated by the success of chimeric antigen receptor (CAR) T cells for hematological malignancies^1,2^ and expanded tumor-infiltrating lymphocytes (TILs) for advanced melanoma ^3,4^. Natural Killer (NK) and γδ T cells are now also being extensively explored as potential immunotherapies^5,6^. While both utilization and impact of cellular therapies are expected to increase in the future due to these positive indications, challenges persist in predicting their potency and improving treatment efficacy.

Our understanding of what drives clinical responses to ACT has evolved over time, in TILs progressing from initial studies highlighting increased numbers of CD8+ T-cells in infusion products^7^ to more nuanced recent studies emphasizing the importance of neoantigen-specific^8^ and stem-like^9^ CD8+ T-cells instead. Together with the incredible success of CAR-T clinical trials^10,11^, where much fewer cells are administered, these findings suggest that superior ACT products could be generated by only expanding cells with a highly potent anti-tumor phenotype.

Given that ACT products are comprised of effector cells considered to be primary mediators of direct tumor killing, a logical assumption is that highly potent cells will react strongly against their targets and could form an effective ACT product on their own. Such potential is often measured via functional responses to target cells, e.g., cytokine secretion or cytotoxic ability. However, these efforts are dominated by bulk population assays which can often obscure these critical populations of interest. Single-cell RNA sequencing has alleviated this issue to some degree; however, its utility is limited by cost, frequent lack of RNA-to-protein-to-function translatability^12^, and inability to select cells of interest for further assays or clinical use. Selection of viable effector cells for further applications is currently achievable via fluorescence and magnetic-activated cell sorting, but these methodologies are limited to using surface molecules as surrogates of cell activation or secretome-capture methods affected by cytokine crosstalk. Accurate identification and selection of viable, potent effector cells with demonstrated killing potential is therefore currently difficult, restricting subsequent downstream study of their phenotypes and generation of selectively expanded ACT products.

The Xdrop double-emulsion droplet technology highlighted here offers a solution by providing a high-throughput platform for single-cell study of effector cell functions, such as direct tumor killing, that is compatible with downstream analysis and expansion of viable cells post-assay.

## The Xdrop technology

Few technologies can accommodate single-cell assessment of cytotoxicity or cytokine secretion^13–17^, often requiring extensive microfluidics know-how and specialized equipment. Furthermore, viable cell populations are then difficult to retrieve in a high-throughput manner^13,14^. The Xdrop instrument is to our knowledge the first commercially available hardware addressing both issues. The double-emulsion droplet technology facilitates high throughput compartmentalization of viable single cells or cell pairs in droplets, allowing the study of individual cells or cell-cell interactions over time (**Figure 1**). The technology uses the same droplet encapsulation technology previously employed for microbes and DNA^18,19^, instead utilizing a larger 50 μm diameter double-emulsion droplet (DE50) to accommodate eukaryotic cells.

**Figure 1.**
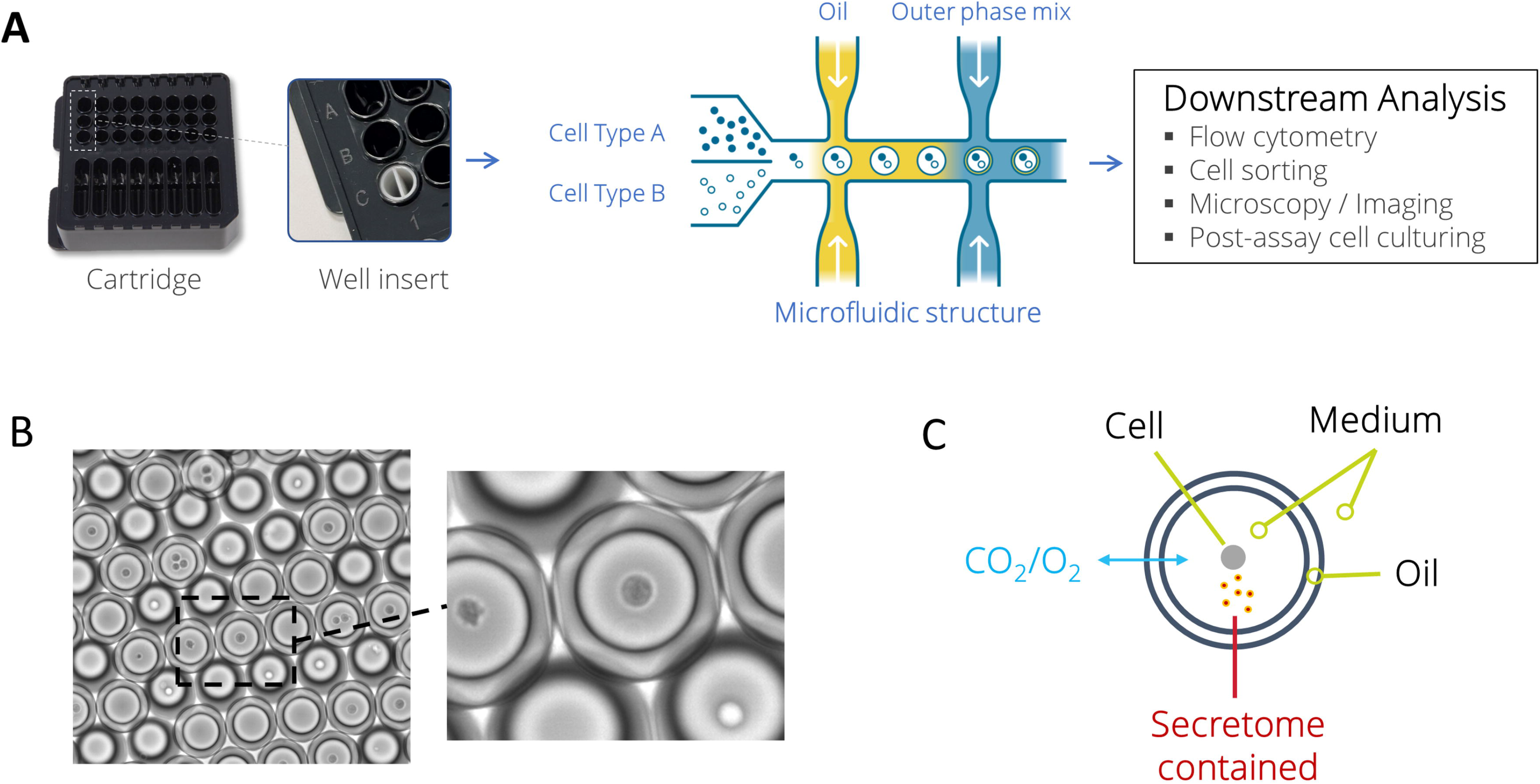
Overview of DE50 droplet generation and characteristics. **(A)** Workflow for DE50 droplet generation and potential analyses. Microfluidic DE50 cartridges are loaded with oil, outer phase mix, and cells, covered with a gasket, and placed into the Xdrop instrument. Up to eight productions can be run in parallel per cartridge, with a run time of ∼8 minutes. When using two different cell types, the Xdrop Well Insert ensures separation of effector and target cells until DE50 droplet encapsulation. Additional assay reagents are added to the cell suspension according to the user’s experimental needs. DE50 droplets are harvested from the cartridge and incubated as necessary whilst the functional assay of choice occurs individually within each droplet. Downstream analysis can be conducted at various time points with a range of potential readouts. **(B)** Microscopy image of DE50 droplets containing cells. **(C)** Schematic diagram of DE50 droplet structure, including passive diffusion of smaller molecules to sustain viability and the retention of the secretome and cell/cells within the droplet.

Prior to encapsulation, cells are stained with fluorescent cell-labelling dyes to facilitate deconvolution during analysis. During the production process, DE50 droplets can be loaded with standard cell culture media, varied cell types, and supplementary reagents (e.g., viability stains and stimulatory molecules) tailored to specific readout requirements. The droplets are stable immediately post-production and can be transferred to common plastic laboratory cell culturing containers equipment using routine pipetting. To preserve cellular viability, droplets are maintained in standard CO_2_ incubators. Analysis of droplets can then be conducted at any timepoint of interest using conventional cell analysis methods such as microscopy and flow cytometry. Continued culturing of cells post-assay is possible by treating sorted droplets with a break reagent, similar to the Xdrop DE20 targeted enrichment procedure for recovery of DNA^18,19^.

## Enrichment of potent cytokine secretors

Cytokine secretion upon stimulation is commonly detected within culture medium through singleplex (e.g., ELISA, ELISpot) or more elaborate multiplexed approaches (e.g., bead-based immunoassays). However, these assays do not permit single-cell deconvolution. An alternative is intracellular cytokine staining which provides single-cell resolution, yet by nature measures intracellular accumulation of cytokines rather than actual secretion. Importantly, none of these approaches allow the viable isolation of specific cells.

Contemporary detection technologies have evolved to capture secreted cytokines on the surface of the secreting cell without impacting viability, potentially providing a solution to the described challenges^20^. However, such approaches are vulnerable to cytokine crosstalk, where an excess of produced cytokine or premature saturation of capture reagents can cause false positive and high background signals. Instead, such crosstalk can be eliminated by combining this methodology with single-cell containing DE50 droplets. For example, in PBMC-derived T-cells stimulated to produce cytokines, a distinct high-TNF-α producing subpopulation previously masked in the bulk assay becomes identifiable^21^ (**Figure 2A)** through the use of DE50 droplets, highlighting the importance of such single-cell resolution.

**Figure 2.**
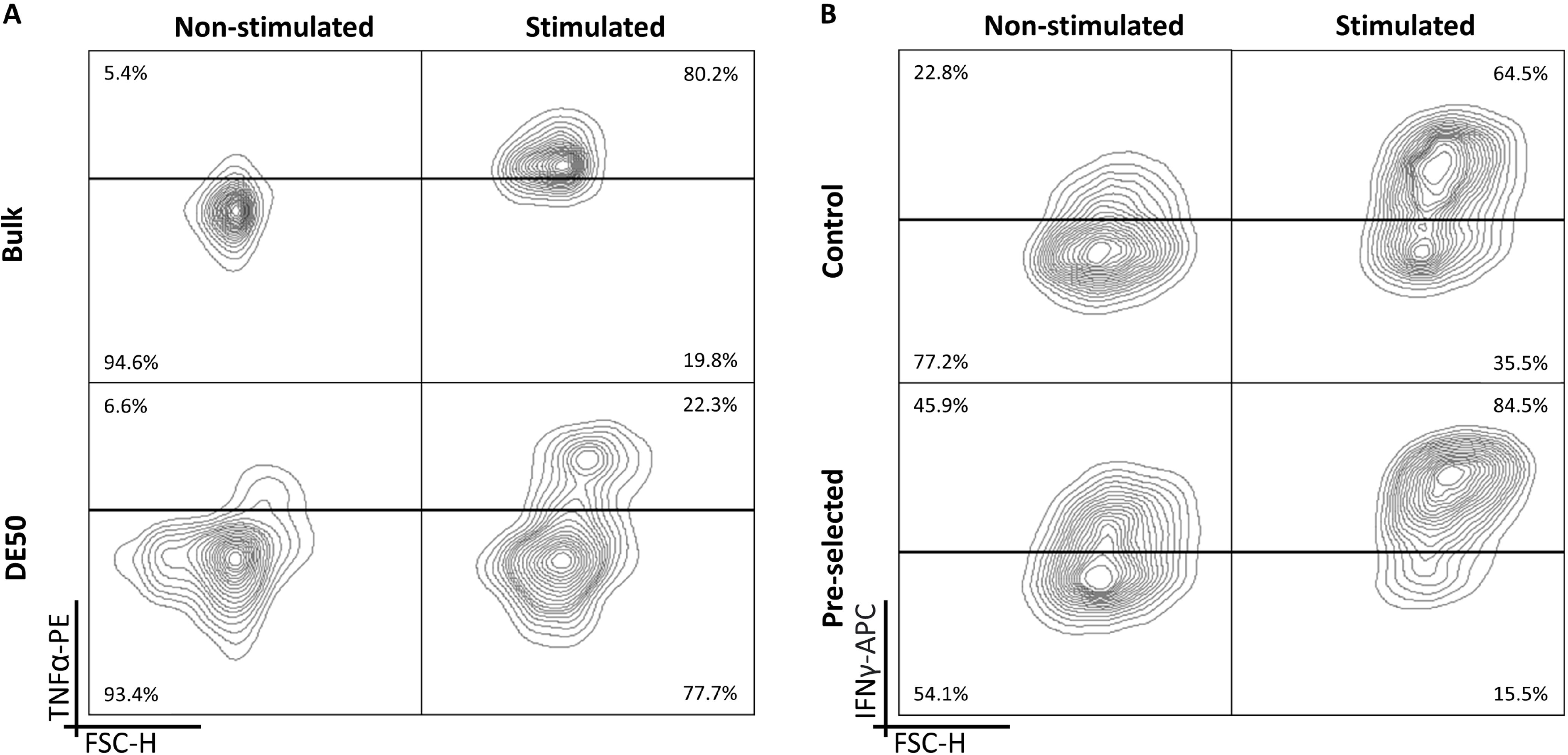
Detection and viable isolation of potent cytokine secreting effector cells using DE50 droplets. **(A)** PBMCs were labelled with TNF-α capture reagents before being split into four groups: bulk samples (+/-PMA/Ionomycin) and DE50 encapsulated samples (+/-PMA/Ionomycin). All groups were then labelled with a TNF-α capture reagent and incubated for four hours. For DE50 samples the TNF-α detection reagent was added during encapsulation whereas for bulk samples the detection reagent was added after four hours. Post-incubation, cells from DE50 droplets were released using droplet break reagent and all samples were stained with α-CD3-PerCP and assessed by flow cytometry. The added resolution from the DE50 encapsulation reveals a high TNF-α-producing T-cell subpopulation not detectable when performing the assay in bulk^21^. **(B)** NK cells were stimulated with IL-2 and labelled with IFN-γ capture reagent before being encapsulated in DE50 droplets together with IFN-γ detection reagent. After four hours of incubation all droplets were broken as for (A) and high IFN-γ-secreting cells were enriched (FACS). Cells were then cultured for an additional two weeks, with control non-stimulated cells cultured in parallel. IFN-γ secretion in the DE50 assay was then measured once after stimulation with IL-2 (or no stimulation, control) of both the non-enriched and enriched populations. The pre-selected population retained their enhanced IFN-γ-secreting abilities in comparison to the non-enriched control, confirming the successful isolation and subsequent expansion of effectors with an indicated increased potency^22^.

The translation of cytokine secretion output into downstream applications, such as deeper analysis or preferential expansion of the most potent secretors during ACT product generation, is often limited by technical constraints. As demonstrated with cytokine-stimulated NK cells^22^ (**Figure 2B**), the DE50 technology bridges this gap by facilitating the sorting of viable cytokine secreting populations of interest. Here a potent IFN-γ secreting subpopulation of NK cells was recovered from DE50 droplets and cultured *in vitro* for an additional two weeks. Subsequent analysis revealed that potent IFN-γ secretors retained their superior secretory abilities upon re-stimulation^22^.

## Detection of effective target cell killers

Cytotoxicity assays are rational markers of effector cell potency that reflect the *in vivo* mode of action, a crucial feature for ACT product potency assays as noted in recent proposed FDA guidance documentation^23^. Isotopic chromium release, bioluminescence, impedance, and flow cytometry-based assays are all commonly employed; however, often the observed effect cannot be linked to specific individual cells. The observed cytotoxic abilities may therefore be due to confounding factors such as highly potent serially killing effectors^24^ masking a lack of cytotoxicity in the bulk of the population. Instead, DE50 droplets provide the single-cell resolution required to appropriately assess the cytotoxic potential of ACT products.

To quantify and identify effective killers using DE50 droplets, target and effector cells are labelled with distinct dyes prior to encapsulation and dead cell-staining propidium iodide (PI) is added immediately prior to encapsulation. The use of these dyes enables clear identification of droplet contents and the real-time labelling of newly dead cells throughout the incubation **(Figure 3A-B)**. Background cell death can be readily estimated within each unique droplet composition and is used to calculate true target-cell killing. Importantly, this single-cell level cytotoxicity assay is compatible with a wide range of ACT-relevant cell types, including NK cells^25^, TILs, and CAR-T cells (**Figure 3C-E**).

**Figure 3.**
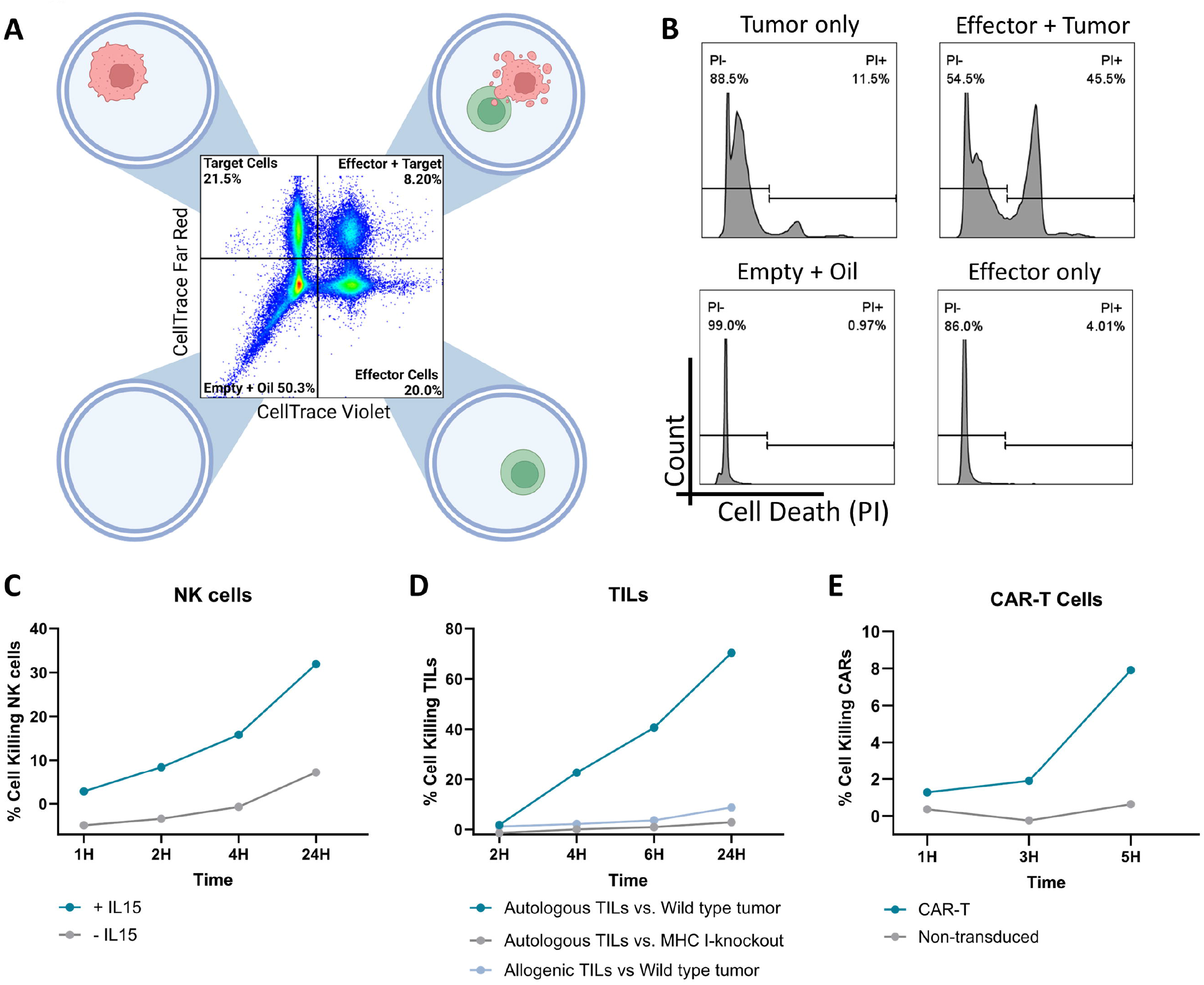
Cross-cell type compatibility of DE50 droplets with propidium-iodide based cytotoxicity assays. **(A)** Distinct fluorescent labelling of effector and target cells prior to encapsulation in DE50 droplets allows deconvolution of droplet composition: empty, target+, effector+, and target+effector+. **(B)** Addition of propidium iodide (PI) to the encapsulation process allows detection of cell death as it occurs within droplets, which can be quantified within each unique sub-population (those defined in A) during analysis to determine background cell death and true effector-induced target cell death. **(C-E)** Representative plots of effector cell-mediated killing of target cells in the described cytotoxicity assay, adjusted for estimated background cell death. **C)** NK cells +/-IL-15 vs K562 target cells^25^. **D)** Autologous/allogeneic TILs vs. autologous wildtype (WT) or MHCI-deficient (*B2M*^-^) tumour. **E)** CD19-specific CAR-T^27^ or non-transduced T-cells vs. CD19+ Daudi target cells. Cytotoxicity at each time point was calculated by normalizing the observed death in co-encapsulating droplets (minus background) to all co-encapsulating droplets (minus background): Percentage of cell killing effector cells at _t(x)_ = (Observed death_t(x)_ –Background death_t(x)_)/(100-Background death_t(x)_). Here background is estimated based on cell death in non-co-encapsulating populations. Background death_t(x)_ = (1-(1-%TargetCellDeath_t(x)_/100)*(1-%EffectorCellDeath_t(x)_/100))*100.

Alternative readouts of cytotoxicity can also be employed in the DE50 droplet setup. Granzyme B (GzmB) is often used as a surrogate marker for cytotoxic ability, and the addition of a cleavage-triggered fluorescent GzmB substrate to the encapsulation process facilitates the detection of its release^26^, as exemplified with CAR-T cells **(Figure 4A-C)**. The granularity of this assay, and subsequent identification of highly potent effectors, can be achieved by combining it with the PI-based cytotoxicity assay^26^. Using CAR-T cells, we demonstrate the simultaneous detection of GzmB secretion and cell killing in DE50 droplets **(Figure 4D-F)**. This combination revealed that not all GzmB+ encapsulations resulted in rapid target-cell killing, emphasizing the value of integrating multiple parameters for enhanced single-cell insights.

**Figure 4.**
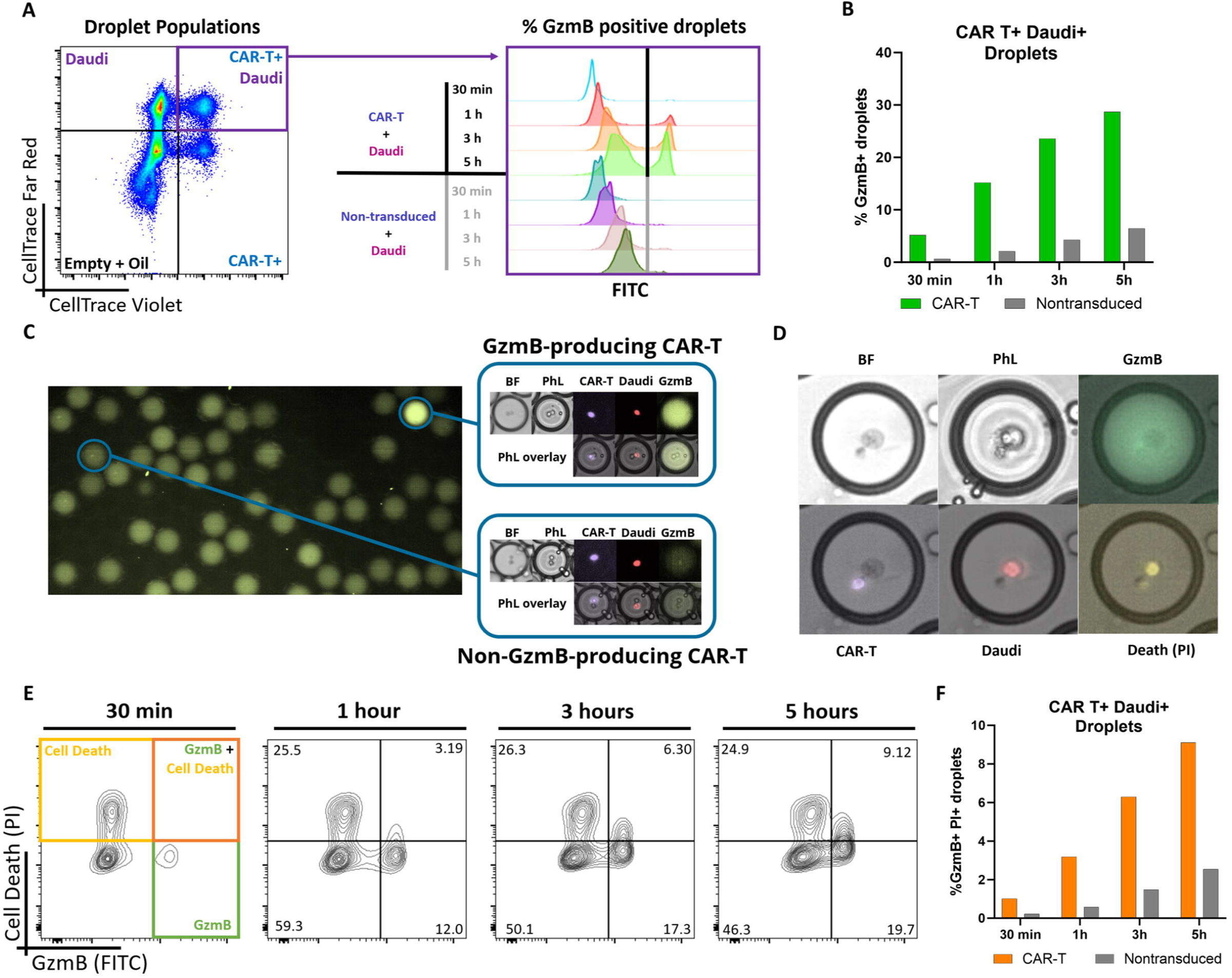
Identification of potent effector cells via detection of granzyme B secretion. **(A-B)** The addition of a fluorescent-tagged granzyme B (GzmB) substrate, which fluoresces upon cleavage of the substrate, during the encapsulation process allows detection of GzmB secretion within DE50 droplets^26^, as demonstrated with CD19-specific CAR-T^27^ and non-transduced T-cells vs CD19^+^ Daudi target cells. **(C)** Representative microscopy images of CAR-T^+^Daudi^+^ DE50 droplets demonstrating the presence, or lack, of GzmB production. **(D-F)** The GzmB assay can be combined with the PI-based cytotoxicity assay^26^ described in Figure 3 to provide additional functional information regarding the cell-killing ability of effector cells, as shown with the same effector/target pairs from (A-C).

## Conclusion and Future Perspectives

The double-emulsion droplet technology presents a robust single-cell level platform that overcomes current limitations in detecting and isolating highly potent effector cells within ACT products. This technology eliminates the masking effects of conventional bulk approaches whilst retaining the viability of cells post-assay.

The ability to isolate viable truly potent effectors allows a wide range of downstream in-depth characterizations concerning general phenotype, proliferative ability, response to repeated stimulation, functional profiles, and other such attributes. The highly customizable nature of the assay promotes flexibility across many cell types and potency parameters, and findings resulting from these kinds of studies are likely to inform the future development of improved ACT by guiding the production of highly potent infusion products. Revealing the true functional profiles of individual cells in ACT products may also produce advances in linking clinical responses to these traits, and in the future, there is substantial potential for direct downstream therapeutic ACT applications. Beyond the identification of highly potent effectors, the technology has potential uses in complementary investigations. For example, is the lack of rapid target cell death detected in many GzmB^+^ droplets **(Figure 4)** caused by poorly functioning T-cells or by target cell resistance mechanisms? Further analysis of the droplets in question would provide some clues.

Although promising, the described technology comes with limitations. Droplets with >2 cells can occur, but this issue can be mitigated by optimizing loading ratios and concentrations. Similarly, optimizing pre-assay cell handling procedures reduces background cell death signals, which are likely cell-type specific. A small degree of background cell death still occurs, but this can easily be accounted for when calculating target cell killing. The cytotoxicity assays presented here cannot definitively confirm which cell of the pair within the droplet has died when analyzed using flow cytometry; however, complementary imaging analyses can confirm the observed/expected trends, as demonstrated in Figure 4D. Alternatively, target cell lines engineered to express specific fluorescent markers (e.g., GFP/YFP/RFP) upon death are likely an effective strategy for clarifying any ambiguity. Continued expansion post-assay may be affected by the break reagent used to disrupt droplets as it can have cytotoxic effects. This may be especially problematic in the case of extremely rare populations, although effector cell expansion protocols often result in significant fold expansions from even minimal starting material.

In conclusion, the double-emulsion droplet technology has the potential to play a sizable role in properly understanding and harnessing highly potent effector cells in the ACT context.

## Acknowledgments

The authors thank all patients who donated clinical material to this study. Written informed consent was provided by all patients prior to sample collection. Samples were collected from patients enrolled in clinical protocols at the National Center for Cancer Immune Therapy (CCIT-DK), Department of Oncology, Copenhagen University Hospital, Herlev, Denmark. All procedures were approved by the Ethics Committee of the Capital Region of Denmark and national regulations for biomedical research (Ethical approval reference: H-20070020; Data Protection approval P-2021-303). The CD19-specific CAR-T cell construct^27^ was generously provided by the Holt group at the BC Cancer Research Institute (Vancouver, Canada). Biorender.com, GraphPad Prism v10 (GraphPad Software, Boston, Massachusetts, USA), and FlowJo™ v10.0 Software (BD Life Sciences, Ashland, Oregon, USA) were used for figure generation. This research was supported by a grant from the European Innovation Council Project 190144395 to Samplix ApS. Samplix® and Xdrop® are registered trademarks of Samplix ApS.

## Declaration of Generative AI and AI-assisted technologies in the writing process

During the preparation of this work, the authors used ChatGPT3.5 and Grammarly v1.2 in order to improve the readability of the manuscript. After using these tools, the authors reviewed and edited the content as needed and take full responsibility for the content of the publication.

